# A dual gene-specific mutator system installs all transition mutations at similar rates *in vivo*

**DOI:** 10.1101/2022.06.09.495438

**Authors:** Daeje Seo, Ga-eul Eom, Hye Won Kim, Seokhee Kim

## Abstract

Targeted *in vivo* hypermutation accelerates directed evolution of proteins through concurrent DNA diversification and selection. Among recently developed methods, the systems employing a fusion protein of a nucleobase deaminase and T7 RNA polymerase present gene-specific targeting. However, their mutational spectra have been largely limited to exclusive or dominant C:G→T:A mutations. Here we describe eMutaT7^transition^, a new gene-specific mutator system, that installs all the transition mutations (C:G→T:A and A:T→G:C) at comparable rates. By using two mutator proteins in which two efficient deaminases, PmCDA1 and TadA-8e, are separately fused to T7 RNA polymerase, we obtained similar numbers of C:G→T:A and A:T→G:C mutations at a sufficiently high rate (∼3.4 × 10^-5^ mutations per base per generation or ∼1.3 mutations per 1 kb per day). Through eMutaT7^transition^-mediated TEM-1 evolution for antibiotic resistance, we generated many mutations also found in clinical isolates. Overall, with a fast mutation rate and wider mutational spectrum, eMutaT7^transition^ is a potential first-line method for gene-specific *in vivo* hypermutation.

## INTRODUCTION

Directed evolution is a powerful approach that mimics natural evolution to improve biomolecular activity (Arnold, 1998; Packer & Liu, 2015). Traditional directed evolution relies on *in vitro* gene diversification such as error-prone PCR or randomized oligonucleotide pools (Packer & Liu, 2015). In contrast, continuous directed evolution (CDE) adopts *in vivo* hypermutation, allowing simultaneous gene diversification, selection, and replication in cells; this technique significantly enhances the depth and scale of biomolecular evolution (Molina *et al*, 2022; Morrison *et al*, 2020; Rix & Liu, 2021). As random mutagenesis in the genome is highly deleterious to cells, *in vivo* hypermutation methods should aim to introduce mutations in a relatively narrow region around the target gene (Molina *et al*., 2022).

Various methods for targeted *in vivo* hypermutation have been reported recently, based distinct molecular principles and thus presenting assorted target ranges (Molina *et al*., 2022; Morrison *et al*., 2020; Rix & Liu, 2021). Several CRISPR-Cas-based methods (e.g., EvolvR (Halperin *et al*, 2018), CRISPR-X (Hess *et al*, 2016), base editors (Komor *et al*, 2016; Ma *et al*, 2016; Nishida *et al*, 2016), and prime editors (Anzalone *et al*, 2019)) install mutations in smaller regions of a gene. OrthoRep (Ravikumar *et al*, 2018) uses an orthogonal error-prone DNA polymerase and generates mutations on a plasmid. Virus-based methods, such as PACE (Esvelt *et al*, 2011) and mammalian cell-based systems (Berman *et al*, 2018; English *et al*, 2019), mutate the entire viral genome. Considering that most directed evolution experiments focus on a single protein, the ideal target range is a single gene of interest. Gene-specific targeting was achieved through homologous platforms that use chimeric mutator proteins, generated by fusing a nucleobase deaminase to an orthogonal T7 RNA polymerase (T7RNAP) (Alvarez *et al*, 2020; Butt *et al*, 2021; Chen *et al*, 2020; Cravens *et al*, 2021; Moore *et al*, 2018; Park & Kim, 2021).

The deaminase-T7RNAP system was first reported in bacteria (MutaT7; (Moore *et al*., 2018)) and further extended to mammalian cells (TRACE; (Chen *et al*., 2020)), yeast (TRIDENT; (Cravens *et al*., 2021)), and plants (Butt *et al*., 2021). We previously demonstrated that the mutation rate of MutaT7 could be enhanced 7-to 20-fold with a more efficient cytidine deaminase, *Petromyzon marinus* cytidine deaminase (PmCDA1) (Park & Kim, 2021). This PmCDA1_T7RNAP mutator (previously termed eMutaT7, but here renamed eMutaT7^PmCDA1^) generated ∼4 mutations per 1 kb per day (∼9.4 × 10^-5^ mutations per base per generation) in *Escherichia coli*, representing the fastest gene-specific *in vivo* mutagenesis. The major limitation of eMutaT7^PmCDA1^ is a narrow mutational spectrum: it mainly generates C→T mutations on the coding strand and, with the Shoulders group’s dual promoter/terminator approach that allows transcription in both directions, introduces C→T and G→A mutations (C:G→T:A) (Moore *et al*., 2018; Park & Kim, 2021). Mutations could be expanded to A→G and T→C (A:T→G:C) with engineered tRNA adenosine deaminases, TadA-7.10 (Alvarez *et al*., 2020; Gaudelli *et al*, 2017) and yeTadA1.0 (Cravens *et al*., 2021), but they either had a ∼200-fold slower mutation rate (∼5 × 10^-7^ mutations per base per generation) (Alvarez *et al*., 2020) than eMutaT7^PmCDA1^, or presented C:G→T:A as dominant mutations (∼95%) in nonselective conditions when combined with PmCDA1_T7RNAP (Cravens *et al*., 2021).

Here, we report on eMutaT7^transition^, a new dual mutator system that introduces all transition mutations (C:G→T:A and A:T→G:C) at comparable rates. The eMutaT7^transition^ system uses two mutators, eMutaT7^PmCDA1^ and eMutaT7^TadA-8e^. The latter is the fusion of T7RNAP and a recently evolved *E. coli* adenosine deaminase, TadA-8e (Richter *et al*, 2020), which had much higher mutational activity than the previously evolved TadA-7.10 (Gaudelli *et al*., 2017) (Fig 1). We optimized the expression of the two mutators and a uracil glycosylase inhibitor, and demonstrated that the frequencies of the C:G→T:A and A:T→G:C mutations were not significantly different. Furthermore, overall mutation rate was not markedly reduced. eMutaT7^transition^ also promoted rapid continuous directed evolution of antibiotic resistance with various transition mutations, suggesting that it is a viable alternative for gene-specific *in vivo* hypermutation with an improved mutational spectrum.

**Figure 1.**
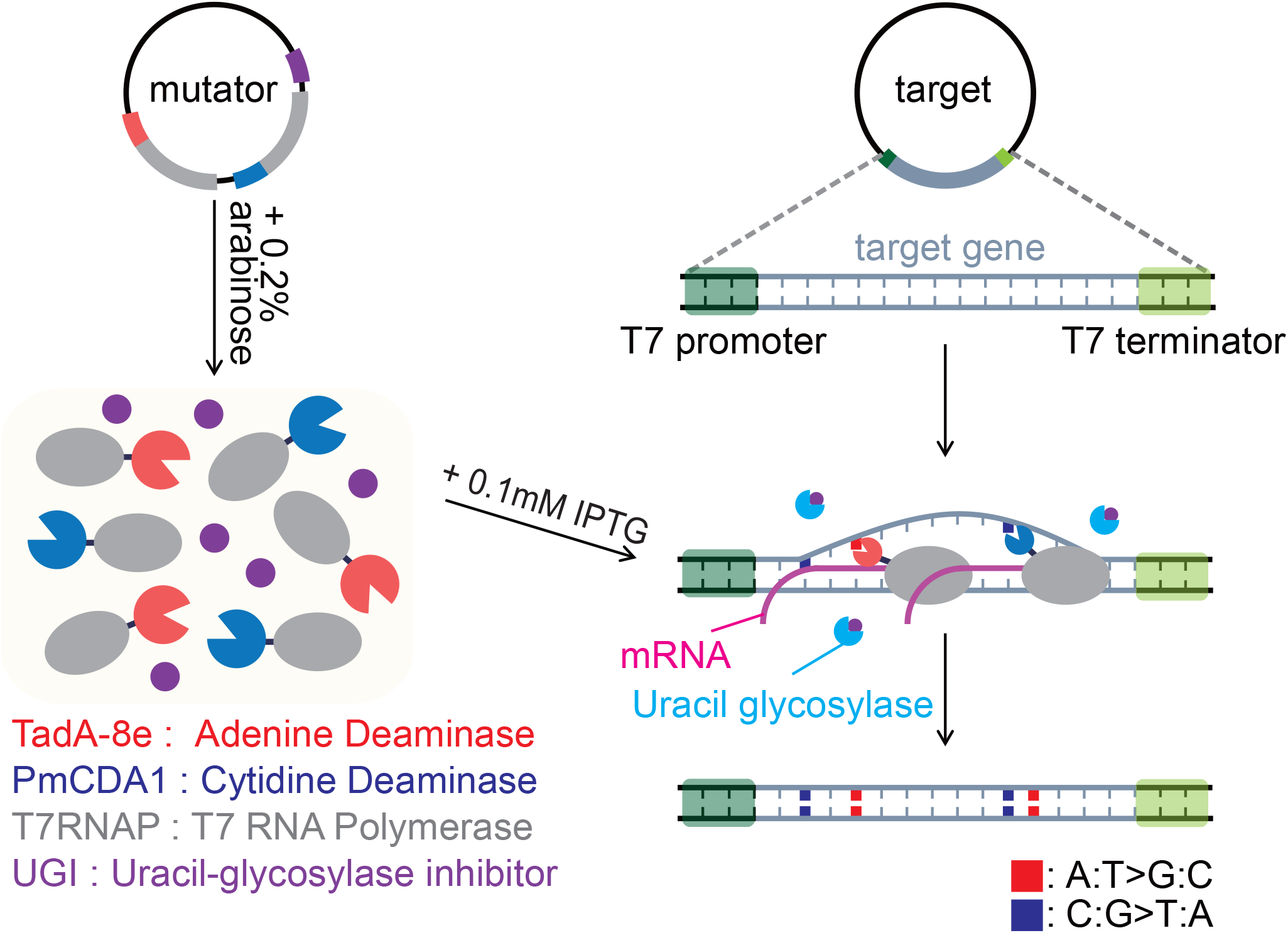
Schematic of the eMutaT7^transition^ system. eMutaT7^transition^ uses two chimeric mutators, eMutaT7^PmCDA1^ and eMutaT7^TadA-8e^, and co-expresses a uracil-glycosylase inhibitor (UGI). The two mutators are expressed with 0.2% arabinose, and target gene transcription is induced with 0.1 mM IPTG. When T7 RNA polymerases transcribe the target gene between T7 promoter and T7 terminator, deaminase enzymes introduce C:G→T:A (cytidine deaminase) or A:T→G:C (adenine deaminase) mutations. UGI promotes C:G→T:A mutations via preventing uracil removal in DNA (Wang *et al*, 1991).

## RESULTS AND DISCUSSION

### eMutaT7^TadA-8e^ promotes rapid gene-specific *in vivo* hypermutation

To date, TadA-8e is the most efficient TadA variant, presenting a rate constant (*k*_app_) 590 times higher than that of the previous TadA-7.10, and has been successfully used for genome editing (Richter *et al*., 2020). To evaluate their efficiency in gene-specific *in vivo* hypermutation, we fused TadA-8e and TadA-7.10 to the N-terminus of T7RNAP, creating eMutaT7^TadA-8e^ and eMutaT7^TadA-7.10^, respectively (Fig 2A). As in the previous characterization of eMutaT7^PmCDA1^ (Park & Kim, 2021), we expressed the mutator and induced hypermutation in the target gene, *pheS*_A294G, which was inserted between T7 promoter and T7 terminator in a low-copy-number plasmid. We determined mutational suppression of the *pheS*_A294G toxicity by counting viable cells in the presence of *p*-chloro-phenylalanine (*p*-Cl-Phe), which is toxic to cells containing intact *pheS*_A294G. We performed 20 rounds of *in vivo* hypermutation (4 h growth and 100-fold dilution to a new medium for a single round) without *p*-Cl-Phe and then sampled cells at different time points for the cell viability assay (Fig EV1A). We found that eMutaT7^TadA-8e^ suppression frequencies were ∼1000-fold higher than eMutaT7^TadA-7.10^ frequencies after 8 h, indicating that the former induces gene-specific hypermutation much faster than the latter (Fig 2B).

**Figure 2.**
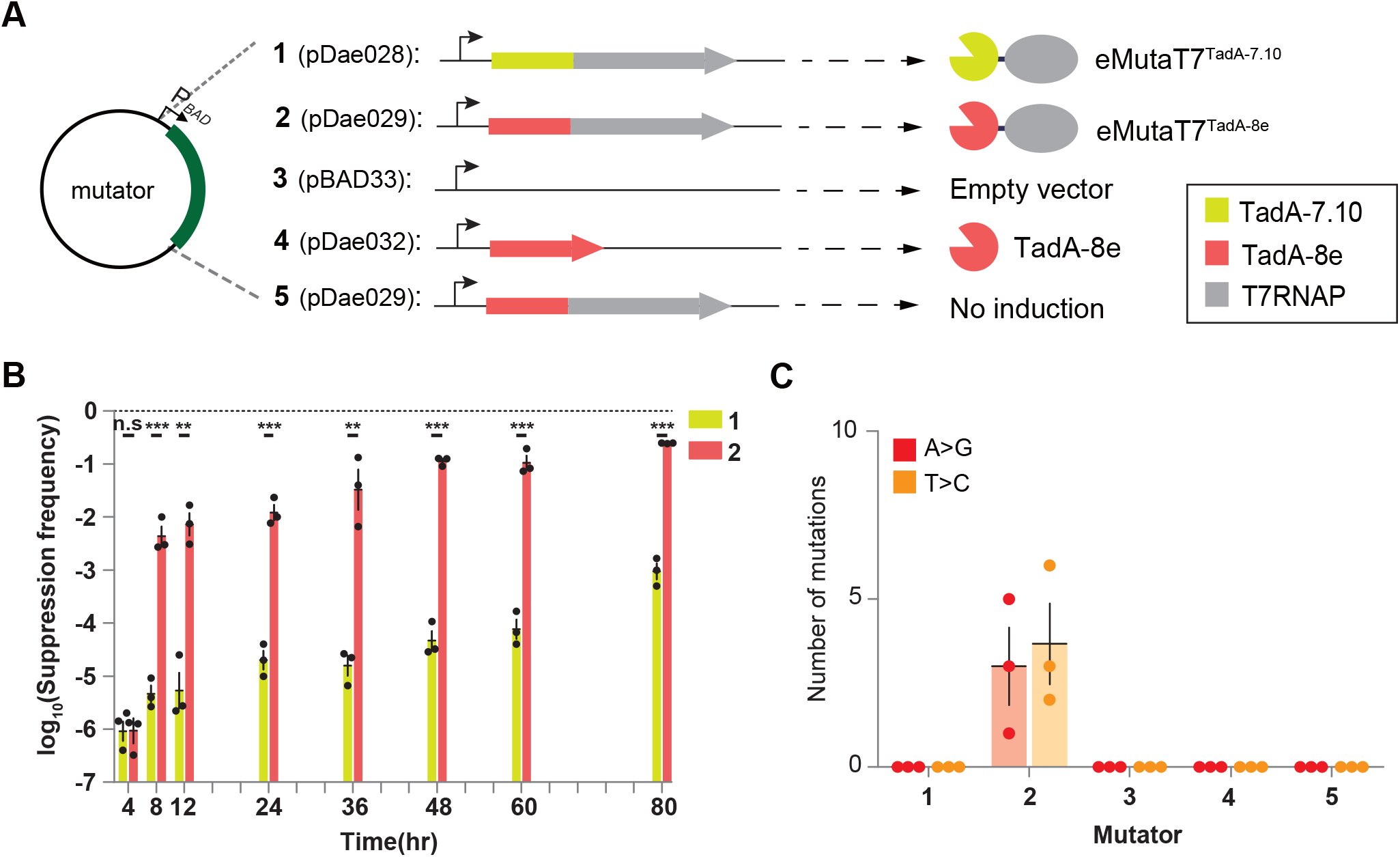
eMutaT7^TadA-8e^ rapidly introduces A→G and T→C mutations in the target gene. A Scheme of the tested mutators and conditions. B Frequency of the *pheS*_A294G toxicity suppression at each mutagenesis cycle for cells expressing eMutaT7^TadA-7.10^ or eMutaT7^TadA-8e^. C Number of A→G (red) or T→C (orange) mutations found in three clones obtained after 20 mutagenesis cycles. Data are presented as dot plots with mean ± standard deviation (SD) (n = 3). \*\**P* < 0.01, \*\*\**P* < 0.001; Student’s *t*-test.

To examine whether eMutaT7^TadA-8e^ generates mutations in the target gene, we randomly selected three clones from cells that had undergone 20 rounds of hypermutation and sequenced the target gene by Sanger method. We also included as negative controls cells that had an empty vector, expressed TadA-8e without T7RNAP, or contained the eMutaT7^TadA-8e^ plasmid without induction (Fig 2A). Notably, we found ∼6.7 mutations per clone (∼4.0 × 10^-5^ mutations per base per generation; ∼1.6 mutations per 1 kb per day) in the eMutaT7^TadA-8e^-expressing cells, while eMutaT7^TadA-7.10^-expressing cells and negative controls did not exhibit mutations (Figs 2C and EV1B). This mutation rate is approximately 80-fold higher than that of eMutaT7^TadA-7.10^ (Alvarez *et al*., 2020) and only 2.4-fold lower than that of eMutaT7^PmCDA1^ (Park & Kim, 2021). Interestingly, we identified nine A→G (45%) and 11 T→C (55%) mutations on the coding strand, indicating that eMutaT7^TadA-8e^ causes mutations on both DNA strands (Figs 2C and EV1B). We observed that eMutaT7^TadA-8e^ neither noticeably reduced cell viability (Fig EV1C) nor induced rifampicin resistance (Fig EV1D). This result suggests that eMutaT7^TadA-8e^ does not generate significant off-target mutations in the genome.

### Deletion of genes associated with hypoxanthine repair does not significantly increase eMutaT7^TadA-8e^ activity

In the eMutaT7^PmCDA1^ system, deletion of a gene encoding a uracil-DNA glycosylase (UNG) enhanced the mutation rate (Park & Kim, 2021). UNG removes uracil (deaminated cytosine) and initiates the base excision repair pathway (Lindahl, 1974). Likewise, we hypothesized that the deletion of genes encoding hypoxanthine (deaminated adenine)-removal enzymes would further increase eMutaT7^TadA-8e^ mutation rate. We prepared a strain in which two genes involved in hypoxanthine repair, *nfi* (Guo *et al*, 1997; Yao *et al*, 1994) and *alkA* (Saparbaev & Laval, 1994), are deleted and analyzed eMutaT7^TadA-8e^-mediated hypermutation (Fig 3A). Twenty rounds of targeted hypermutation revealed that mutations in the *Δnfi ΔalkA* strain did not increase significantly from wild-type levels (average 9 and 6.3 mutations per clone on average, respectively) (Figs 3B and EV2). Because a DNA repair enzyme often reduces mutation rates by more than an order of magnitude (Beletskii & Bhagwat, 1996; Duncan & Weiss, 1982; Foster *et al*, 2015; Schaaper, 1993) and construction of a gene deletion strain requires additional experimental steps, we concluded that the *Δnfi ΔalkA* strain has no obvious advantage over the wild-type strain for eMutaT7^TadA-8e^.

**Figure 3.**
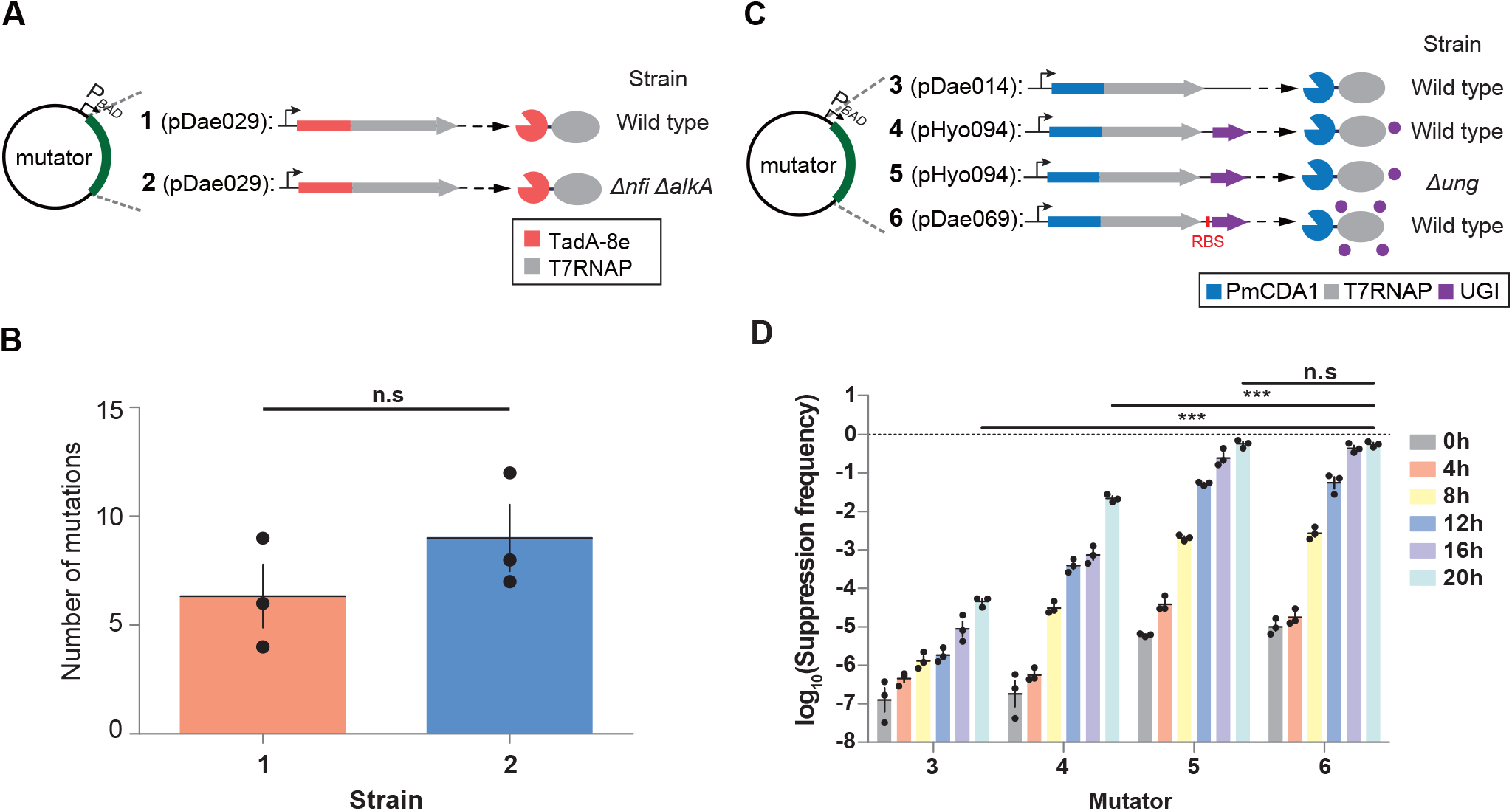
Optimization of eMutaT7^TadA-8e^-or eMutaT7^PmCDA1^-mediated *in vivo* hypermutation. A eMutaT7^TadA-8e^ activity was tested in wild-type or *Δnfi ΔalkA* strains. B Number of mutations found in three clones from the two samples in (A) after 20 mutagenesis cycles. C eMutaT7^PmCDA1^ activity was tested without or with an optimized ribosomal binding site for *ugi* in wild-type or *Δung* strains. D Suppression frequency of the *pheS*_A294G toxicity at each mutagenesis cycle for cells evolved under the four different conditions shown in (C). Data are presented as dot plots with mean ± standard deviation (SD) (n = 3). \*\*\**P* < 0.001, Student’s *t*-test.

### Optimized expression of uracil glycosylase inhibitor increases eMutaT7^PmCDA1^ activity

Although we co-expressed a UNG inhibitor (UGI) with eMutaT7^PmCDA1^ from the plasmid pHyo094, we did not obtain an efficiency level that matched the *Δung* strain (Park & Kim, 2021). Proper UGI expression can greatly expand eMutaT7^PmCDA1^ utility by avoiding the *ung* deletion. To enhance UGI activity, we initially tested a new constitutive promoter for *ugi* or a triply fused protein, UGI_PmCDA1_T7RNAP. However, both were less efficient than the *Δung* strain (Fig EV3). Next, we optimized the ribosomal binding site (RBS) of *ugi* (Salis *et al*, 2009) (Fig 3C), and obtained a suppression frequency indistinguishable from that of the *Δung* strain (Fig 3D). Thus, we were able to avoid *ung* deletion for efficient eMutaT7^PmCDA1^-mediated mutagenesis.

### Dual expression system introduces all transition mutations at comparable rates

We examined whether the two deaminases could simultaneously install both C:G→T:A and A:T→G:C mutations at similar rates. Initially, we tested two triple-fused proteins, PmCDA1_TadA-8e_T7RNAP and TadA-8e_PmCDA1_T7RNAP, in which two deaminases were attached to the N-terminus of T7RNAP in different orders (Fig 4A). Sequencing of clones after 20 rounds of *in vivo* hypermutation revealed that PmCDA1_TadA-8e_T7RNAP installed more A:T→G:C mutations (84%) than C:G→T:A (16%), whereas TadA-8e_PmCDA1_T7RNAP generated more C:G→T:A (96%) than A:T→G:C (4%) (Figs 4B and EV4A). This result indicates that the deaminase closer to T7RNAP is more active. Shorter or longer linker lengths between enzymes did not significantly reduce the gap (Appendix Fig S1).

**Figure 4.**
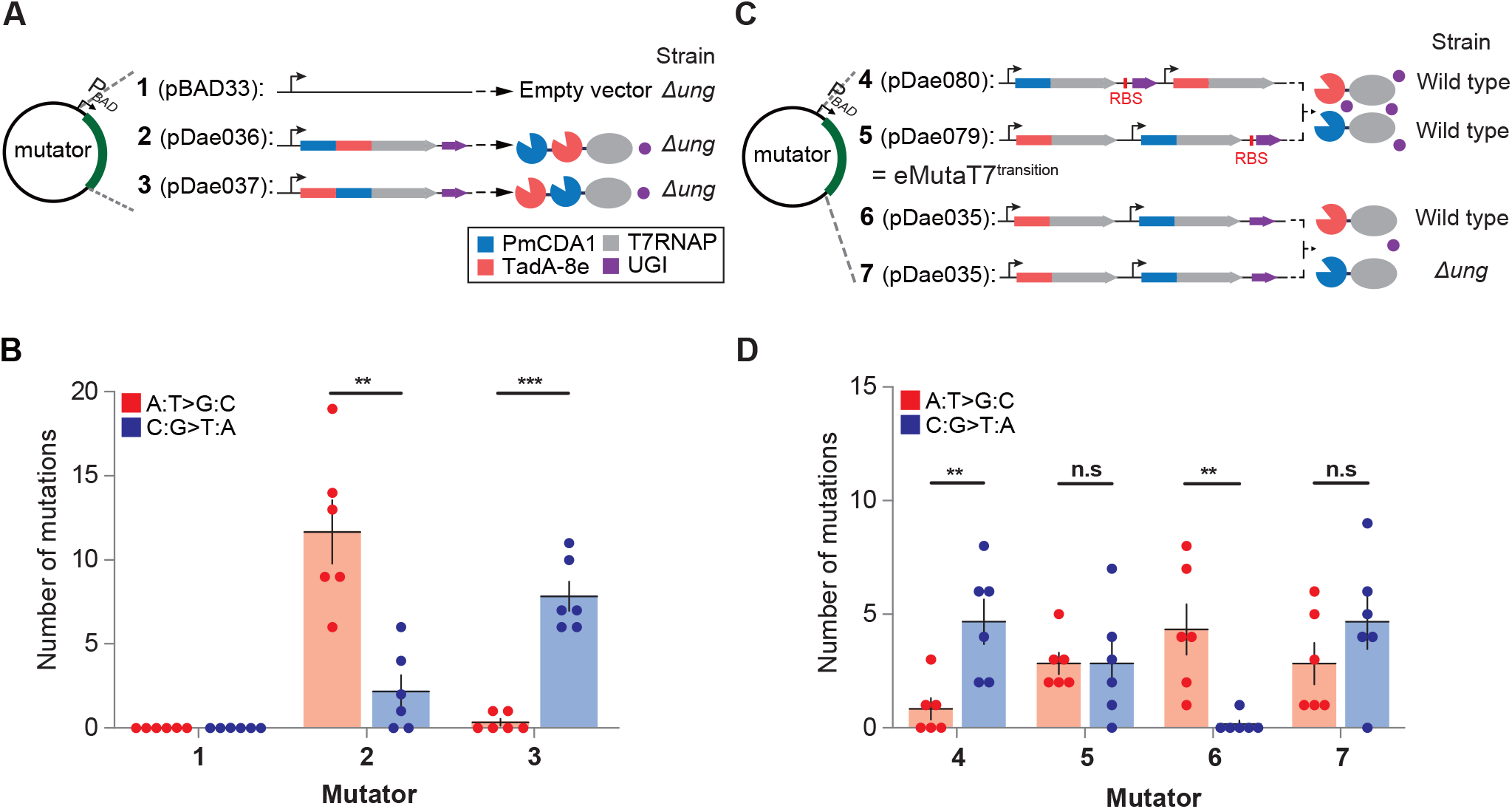
eMutaT7^transition^ rapidly introduces all transition mutations at similar rates. A Two triply-fused mutators were tested for equivalent incorporation of A:T→G:C and C:G→T:A mutations. B Mutation counts in six clones from samples shown in (A) after 20 mutagenesis cycles. C Four dual mutator systems were tested for equivalent incorporation of A:T→G:C and C:G→T:A mutations. D Mutation counts in six clones from samples in (C) after 20 mutagenesis cycles. Data are presented as dot plots with mean ± standard deviation (SD) (n = 6). \*\**P* < 0.01, \*\*\**P* < 0.001; Student’s *t*-test.

Next, we tested the expression of two mutators, eMutaT7^PmCDA1^ and eMutaT7^TadA-8e^, from a single plasmid (Fig 4C). The pDae079 plasmid, in which the eMutaT7^TadA-8e^ gene is located in front of the eMutaT7^PmCDA1^ gene, yielded the same amounts of A:T→G:C (50%) and C:G→T:A (50%) mutations (*p* = 1.0). In contrast, the pDae080 plasmid, which reversed the order of the two mutators, disproportionately generated C:G→T:A (85%) over A:T→G:C (15%) (*p* = 0.0058; Figures 4D and S4B). As expected, weaker UGI expression without optimized RBS significantly reduced C:G→T:A mutations in the wild-type strain (*p* = 0.0041) but produced comparable numbers of mutations in the *Δung* strain (A:T→G:C, 38%; C:G→T:A, 62%; *p* = 0.25; Figs 4D and EV4B). We thus selected pDae079 for eMutaT7^transition^, which appears to install transition mutations at a rate of ∼3.4 × 10^-5^ mutations per base per generation (∼1.3 mutations per 1 kb per day), approximately 2.8-fold slower than that of eMutaT7^PmCDA1^ (Park & Kim, 2021).

### eMutaT7^transition^ evolves TEM-1 with various transition mutations

We previously demonstrated that eMutaT7^PmCDA1^ promoted rapid continuous directed evolution of TEM-1 for resistance against third-generation cephalosporin antibiotics, cefotaxime (CTX) and ceftazidime (CAZ) (Park & Kim, 2021). Here, we tested eMutaT7^transition^ in the same way. We used the dual promoter/terminator approach to install both C→T and G→A mutations (Moore *et al*., 2018; Park & Kim, 2021). By sequentially increasing antibiotic concentrations during multiple rounds of *in vivo* hypermutation, we elevated minimum inhibitory concentrations (MICs) from 0.05 to 400–1600 μg/mL in 80 h for CTX (Fig 5A) and from 0.4 to 4000 μg/mL in 48 h for CAZ (Fig 5B).

**Figure 5.**
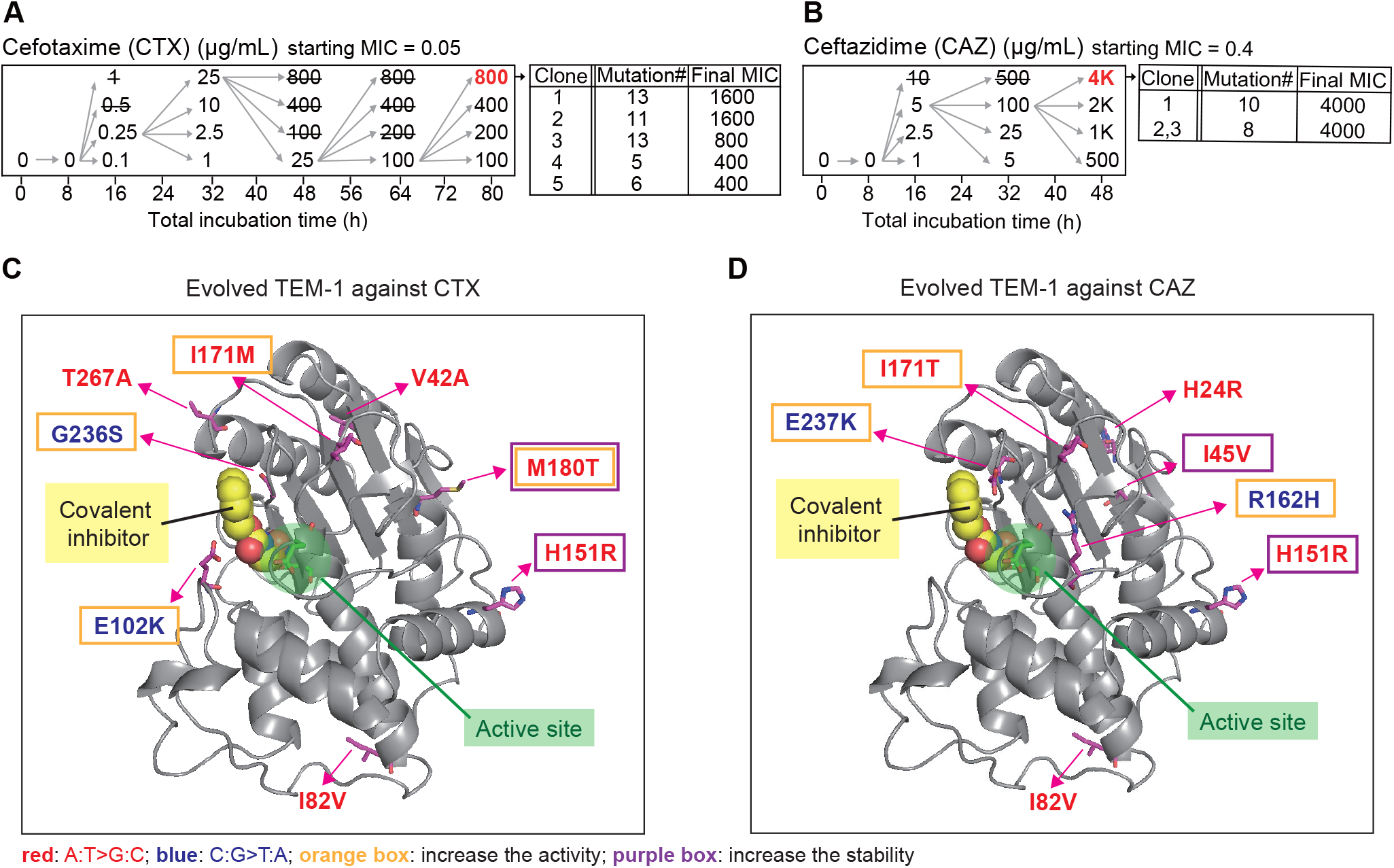
Continuous directed evolution of TEM-1 for antibiotic resistance using eMutaT7transition. A, B Evolutionary pathways of TEM-1 for resistance against CTX (A) or CAZ (B). Each number indicates an antibiotic concentration in a culture. Strikethrough indicates no growth. C, D Structure of TEM-1 (PDB, 1axb) showing a covalent inhibitor (yellow), the active site (green), A:T→G:C mutations in evolved TEM-1 (red), and C:G→T:A mutations in evolved TEM-1 (blue). Mutations in orange and purple boxes indicate those that increased enzyme activity and stability, respectively.

Sanger sequencing of several resistant clones revealed that their mutational spectra were more diverse than those obtained with eMutaT7^PmCDA1^ (Fig EV5). Adenine deamination (A:T→G:C) generated I45V, H151R, I171M, I171T, and M180T, whereas cytosine deamination (C:G→T:A) generated E102K, R162H, G236S, and E237K (Figs 5C, D, and EV5). I45V, H151R, and M180T have been reported to increase enzyme stability (Bershtein *et al*, 2008; Brown *et al*, 2010; Kather *et al*, 2008; Palzkill, 2018). Additionally, E102K, R162H, I171M/T, G236S, and E237K are involved in resistance to CTX or CAZ (Barlow & Hall, 2003; Driffield *et al*, 2006; Palzkill, 2018; Salverda *et al*, 2010; Sowek *et al*, 1991; Zaccolo & Gherardi, 1999). Other mutations are not found in clinical or laboratorial isolates (H24R and V42A), are found in the wild-type allele (I82V), or have unknown functional effect (T267A) (Salverda *et al*., 2010). These results suggest that eMutaT7^transition^ indeed covers a wider protein mutational space for evolution.

In conclusion, this study described a new mutator system that combines eMutaT7^PmCDA1^ and eMutaT7^TadA-8e^, called eMutaT7^transition^. This new system has advantages over previous deaminase-T7RNAP mutators. First, eMutaT7^transition^ expands the mutational spectrum to all transition mutations (C:G→T:A and A:T→G:C). eMutaT7^PmCDA1^ can mediate 8.4% of all amino acid changes (32 out of total 380 changes), but eMutaT7^transition^ expands them to 19% (74 changes). Accordingly, we observed in TEM-1 evolution experiments several A:T→G:C mutations that have been previously identified in clinical or laboratorial isolates. Second, all transition mutations are produced at similar rates. This outcome was made possible by the use of two efficient deaminases, PmCDA1 and TadA-8e, along with appropriate expression of the two mutators and a DNA glycosylase inhibitor. In contrast, TRIDENT generated considerably more C:G→T:A mutations (∼95%) in yeast (Cravens *et al*., 2021).

eMutaT7^transition^ had a ∼2.8-fold slower mutation rate (∼3.4 × 10^-5^ mutations per base per generation; ∼1.3 mutations per 1 kb per day) than eMutaT7^PmCDA1^ (Park & Kim, 2021). However, it was still sufficiently fast to support rapid continuous directed evolution of TEM-1. Moreover, as also observed with eMutaT7^PmCDA1^, evolved TEM-1 variants often contained 2-to 3-fold more mutations than expected from the nominal rate; variants resistant to CTX and CAZ showed rates of ∼2.6 and ∼3.9 mutations per 1 kb per day, respectively (Fig EV5). Future research should aim to include transversion mutations in the mutational spectrum without significantly sacrificing mutation rate. With its good mutation rate and wider mutational spectrum, we believe that eMutaT7^transition^ can become the method of choice for gene-specific *in vivo* hypermutation.

## MATERIALS AND METHODS

### Materials

All PCR experiments were conducted with KOD Plus neo DNA polymerase (Toyobo, Japan). T4 polynucleotide kinase and T4 DNA ligases were purchased from Enzynomics (South Korea). Plasmids and DNA fragments were purified with LaboPass™ plasmid DNA purification kit mini, LaboPass™ PCR purification kit, and LaboPass™ Gel extraction kit (Cosmogenetech, South Korea). Sequences of all DNA constructs in this study were confirmed by Sanger sequencing (Macrogen, South Korea and Bionics, South Korea). Antibiotics (carbenicillin, chloramphenicol, kanamycin, streptomycin, and tetracycline), arabinose, and Isopropyl β-D-1-thiogalactopyranoside (IPTG) were purchased from LPS solution (South Korea). Cefotaxime and ceftazidime were purchased from Tokyo chemical industry (Japan). H-*p*-Chloro-DL-Phe-OH (*p*-Cl-Phe) was purchased from Bachem (Switzerland).

### Plasmid and *E. coli* strain construction

Plasmids, *Escherichia coli* strains, and primers used in this study are listed in Appendix Tables S1-S3, respectively. Genes for adenine deaminase TadA-7.10 (Gaudelli *et al*., 2017) and TadA-8e (Richter *et al*., 2020) were synthesized from Gene Universal (USA). Genes for the *Petromyzon marinus* cytidine deaminase (PmCDA1), XTEN linker, and T7 RNA polymerase (T7RNAP) were amplified from the plasmid expressing eMutaT7 (pHyo094) (Park & Kim, 2021). Genes for adenine deaminases (TadA-7.10 or TadA-8e), linker, and T7RNAP were linked by in vivo assembly (IVA) cloning (Garcia-Nafria *et al*, 2016). Plasmids expressing PmCDA1_XTEN_T7RNAP (eMutaT7^PmCDA1^), TadA-8e_XTEN_T7RNAP (eMutaT7^TadA-8e^), and UGI were cloned by IVA cloning. Also, genes encoding triply-fused proteins, UGI_PmCDA1_T7RNAP, PmCDA1_TadA-8e_T7RNAP, and TadA-8e_PmCDA1_T7RNAP, were also constructed by IVA cloning.

All plasmids expressing variants of mutators or targets (mutation, deletion, and insertion) were constructed using the site-directed mutagenesis PCR method. (Reikofski & Tao, 1992) Plasmids expressing eMutaT7 and UGI in different conditions (deletion of UGI, an optimized ribosomal binding site for UGI, or a constitutive promoter for UGI) were made on pHyo094. Plasmids harboring TadA-8e were made on pDae029. Plasmids expressing PmCDA1_TadA-8e_T7RNAP with different linkers were constructed on pDae036.

For evolution of antibiotic resistance, a target plasmid (pGE158) was constructed from pHyo245, which contains the *pheS*_A294G gene between dual promoter/terminator pairs in a low-copy-number plasmid (Park & Kim, 2021): Ampicillin resistance gene in pHyo245 was replaced with tetracycline resistance gene and *pheS*_A294G was replaced with the *TEM-1* gene by IVA cloning. Tetracycline resistance gene was amplified from the plasmid pREMCM3 (Melancon & Schultz, 2009) and the *TEM-1* gene was obtained from pHyo182 (Park & Kim, 2021).

W3110 *ΔalkA Δnfi* strain (cDJ085) was constructed by homologous recombination method (Sharan *et al*, 2009) The *alkA* and *nfi* genes in W3110 were replaced with the streptomycin resistance gene and the kanamycin resistance gene, respectively. 30 μg/mL of streptomycin or kanamycin was used for selection. Proper gene deletion was confirmed by colony PCR using 2X TOP simple™ DyeMIX-Tenuto (Enzynomics).

### In vivo hypermutation

Three biological replicates of W3110 or the *Δung* strain (cHYO057) harboring a mutator plasmid and a target plasmid (pHyo182 for a single promoter) were grown overnight in LB medium with 35 μg/mL chloramphenicol and 50 μg/mL carbenicillin (cycle #0). On the following day, the overnight cultures were diluted 100-fold in a fresh LB medium supplemented with 35 μg/mL chloramphenicol, 50 μg/mL carbenicillin, 0.2% arabinose, and 0.1 mM IPTG in a 96-deep well plate (Bioneer, South Korea) and incubated at 37°C with shaking (cycle #1). Bacterial cells were diluted every 4 hours and this growth cycle was repeated up to 20 times for accumulation of mutations. At the end of cycle, a fraction of cells were stored at -80°C with 15% glycerol. To identify mutations in the target gene, cells at cycle #20 were streaked on LB-Agar plates with 35 μg/mL chloramphenicol and 50 μg/mL carbenicillin. Three or six colonies were randomly chosen for isolation of target plasmids. The target genes in the purified target plasmids were sequenced by Sanger sequencing. Mutations were counted in the region between 147-bp upstream and 138bp-downstream of the *pheS*_A924G gene (total 1269 bp). Primer 314 and 315 were used for amplification and sequencing of the target gene that has a single promoter system.

### PheS_A294G suppression assay

Suppression frequency of the *pheS*_A294G toxicity was determined as previously described (Park & Kim, 2021). Cells obtained at the endpoint of each cycle (overnight culture for cycle #0) were diluted to OD_600_ ∼ 0.2. Serial 10-fold dilutions of cells (5μl) using LB broth were placed on YEG-agar plates with or without additives (1.6 mM *p*-Cl-Phe, 0.2% arabinose, and 0.1 mM IPTG) and grown overnight at 37°C. On the following day, the number of colonies on each condition was counted to calculate the suppression frequency. The suppression frequency was calculated as N_1_/N_0_ (N_1_: colony forming unit (CFU) in the *p*-Cl-Phe plates and N_0_: CFU in plates without *p*-Cl-Phe).

### Assays for cell viability and off-target mutagenesis

Cell viability and off-target mutagenesis were assayed as previously described (Park & Kim, 2021). Overnight cultures of the cells harboring the plasmid expressing eMutaT7^TadA-8e^, no mutator, or MP6 were diluted 100-fold in LB supplemented with 35 μg/mL chloramphenicol and grown to a log phase (OD_600_ = 0.2-0.5) at 37°C. Cells were diluted to OD_600_ ∼ 0.2 and serial 10-fold dilutions of cells (5μl) using LB broth were placed on LB-agar supplemented with 35 μg/mL chloramphenicol and 0.2% arabinose. After overnight growth at 37°C, the number of colonies on the plates were counted to calculate CFU/mL. To evaluate the off-target mutagenesis via rifampicin resistance, cells taken at cycle #0 and cycle #20 were grown to log phase in LB supplemented with 35 μg/mL chloramphenicol and 50 μg/mL carbenicillin, and subjected to viability assay on plates with or without rifampicin (50 μg/mL).

### Mutation rate calculation

Mutation rates were calculated and presented as mutations per base per generation and mutations per 1 kb per day. The sizes of the gene region are 1269 bases for *pheS*_A294G and 1121 bases for *TEM-1*. One growth cycle, which is 100-fold growth in four hours, indicates 6.64 generations (log_2_100). The mutation rate of TadA-7.10_T7RNAP from Fernandez group’s report (Alvarez *et al*., 2020) was calculated as (3.0 × 10^-4^ variant calling frequency from Figure 4f) / (200 base) / (3 generation) = 5 × 10^-7^ mutations per base per generation.

### TEM-1 evolution and identification of the evolved mutants

TEM-1 evolution experiments were performed as previously described (Park & Kim, 2021). Strains were grown in LB medium supplemented with 6 μg/mL tetracycline, 35 μg/mL chloramphenicol, 0.2% arabinose, and 0.1 mM IPTG. Cells were grown without selection pressure at the initial cycle. Then, multiple cultures were grown with different concentrations of an antibiotic (cefotaxime and ceftazidime) at the same time and the culture grown at the highest antibiotic concentration (OD_600_ > 1) were used for the next round of evolution. After the final cycle, the target plasmids were purified and re-inserted into fresh W3110 cells harboring the T7RNAP-expressing plasmid (pHyo183) for validation of antibiotic resistance. Twelve colonies were randomly selected for MIC measurement and those with high MIC values (five colonies with 400-1600 μg/mL MIC for CTX, three colonies with 4000 μg/mL MIC for CAZ) were subjected to the target gene sequencing by Sanger method.

### MIC determination

MIC values were measured as previously described (Park & Kim, 2021). Cells were grown overnight in LB medium supplemented with 6 μg/mL tetracycline, 35 μg/mL chloramphenicol. They were diluted 10,000-fold into fresh LB broth with increasing concentrations of antibiotics (2-fold) in 96-deep well plates, and grown at 37°C with shaking (290 rpm) overnight. Final cell density (OD_600_) was measured by M200 microplate reader (TECAN, Switzerland).

## ACKNOWLEDGEMENTS

We thank Hyojin Park, Younghyun Kim, Chanwoo Lee, Hyunjin Cho, Inseok Song, Hyunbin Lee, Hyunsung Nam, Kijeong Yang, and Chung-Mo Park for comments and helpful discussions. We are grateful to Nam Ki Lee and Soojung Lee for providing purified DNAs, and Woon Ju Song for providing rifampicin. This work was supported by the National Research Foundation of Korea (NRF) grant funded by the Korea government (MSIT) (2021R1A2C1008730).

## AUTHOR CONTRIBUTIONS

DS and SK developed the initial concept; DS, GE, and HWK performed experiments and analyzed the results; SK supervised the work throughout; DS and SK wrote the manuscript.

## EXPANDED VIEW FIGURE LEGENDS

**Figure EV1.**
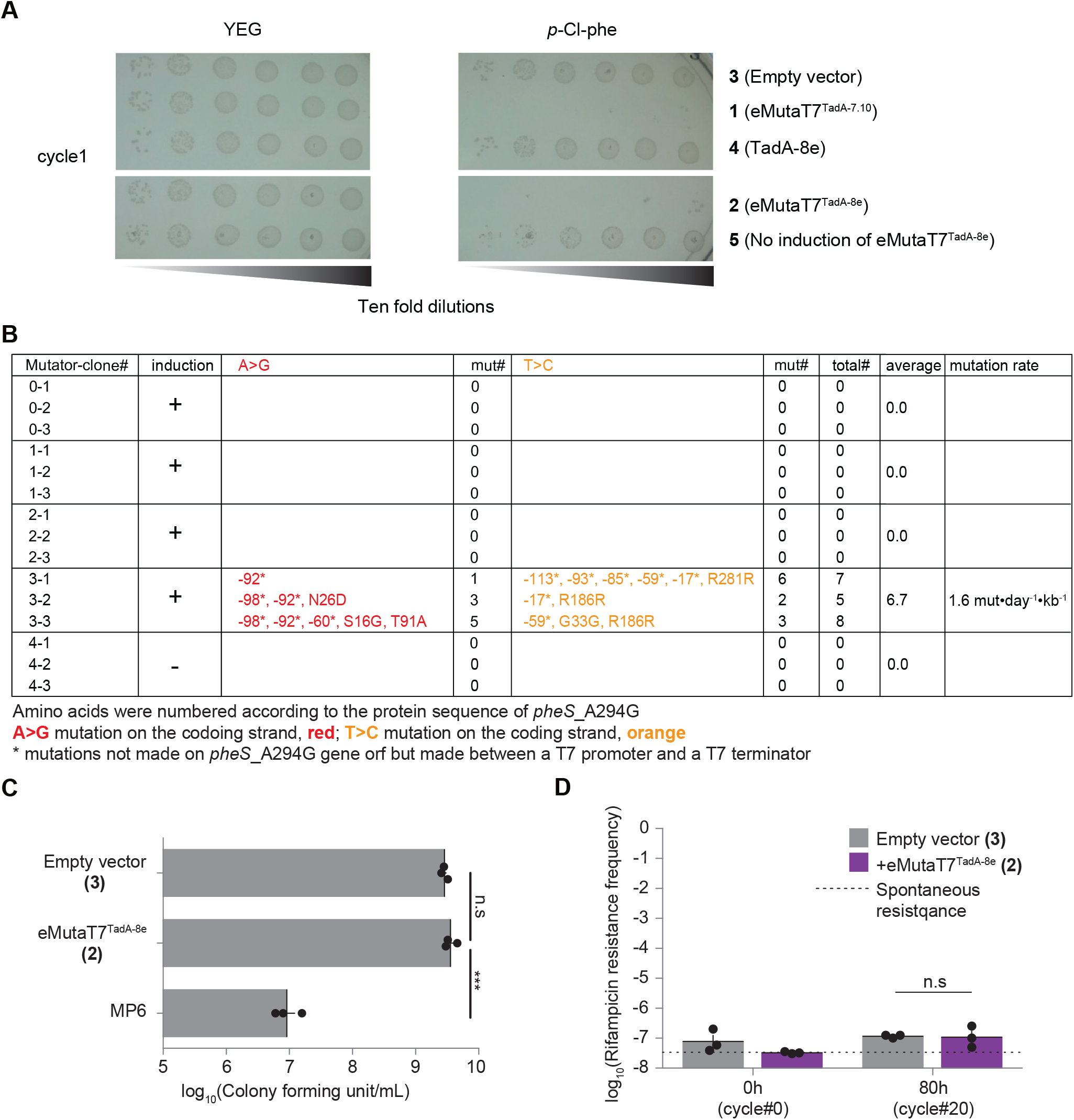
Characterization of eMutaT7^TadA-8e^. A A representative result of the *pheS*_A294G toxicity suppression experiment without or with *p*-Cl-Phe. Strains harboring the *pheS*_A294G target gene were subjected to *in vivo* hypermutation in five different conditions: expressing eMutaT7^TadA-7.10^, eMutaT7^TadA-8e^, no protein, TadA-8e, or no eMutaT7^TadA-8e^ from the cognate plasmid. Cells taken at particular growth cycle were grown in LB media (without arabinose and IPTG) until OD_600_ reached ∼0.2. Serial 10-fold dilutions of cells were spotted and grown on YEG-agar plates supplemented with or without 1.6 mM *p*-Cl-Phe. Growth of cells expressing *pheS*_A294G is inhibited in the presence of *p*-Cl-Phe. Only the eMutaT7^TadA-7.10^-and eMutaT7^TadA-8e^-expressing cells had T7 RNA polymerase and thus showed growth inhibition in the presence of *p*-Cl-Phe. B A list of mutations found in samples shown in Figure 2C. C Viability of cells expressing no protein, eMutaT7^TadA-8e^, or MP6. D Off-target mutation level of cells expressing eMutaT7^TadA-8e^ or no protein was estimated by rifampicin resistance frequency. The dotted line represents spontaneous rifampicin resistance level. Data are presented as dot plots with mean ± standard deviation (SD) (n = 3). ***P < 0.001; Student’s t-test.

**Figure EV2.**
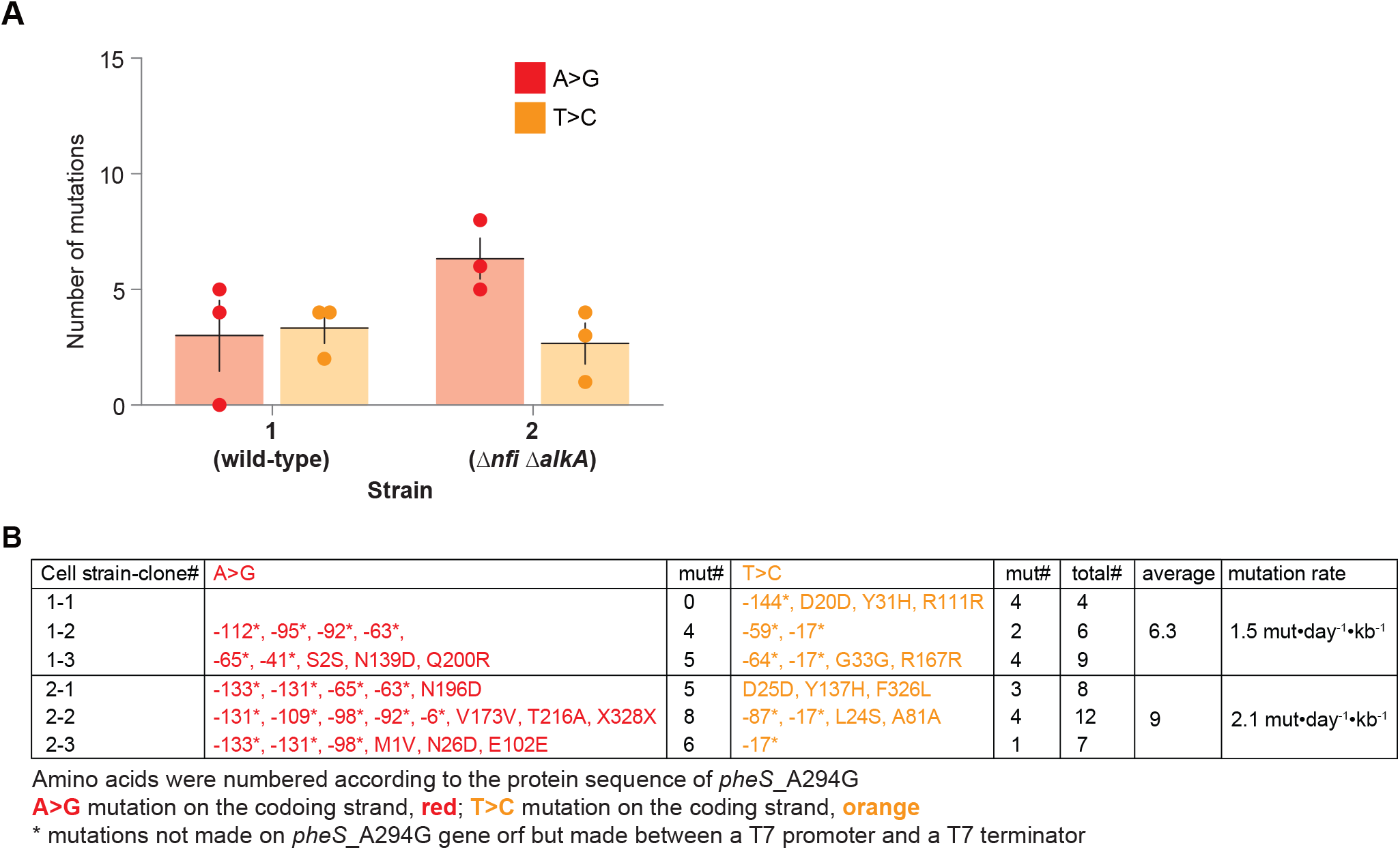
The deletion of inosine glycosylases did not elevate the mutation rate of eMutaT7^TadA-8e^. A Number of A>G (red) and T>C (orange) mutations in samples shown in Figure 3B. B A list of mutations in (A). Data are presented as dot plots with mean ± standard deviation (SD) (n = 3).

**Figure EV3.**
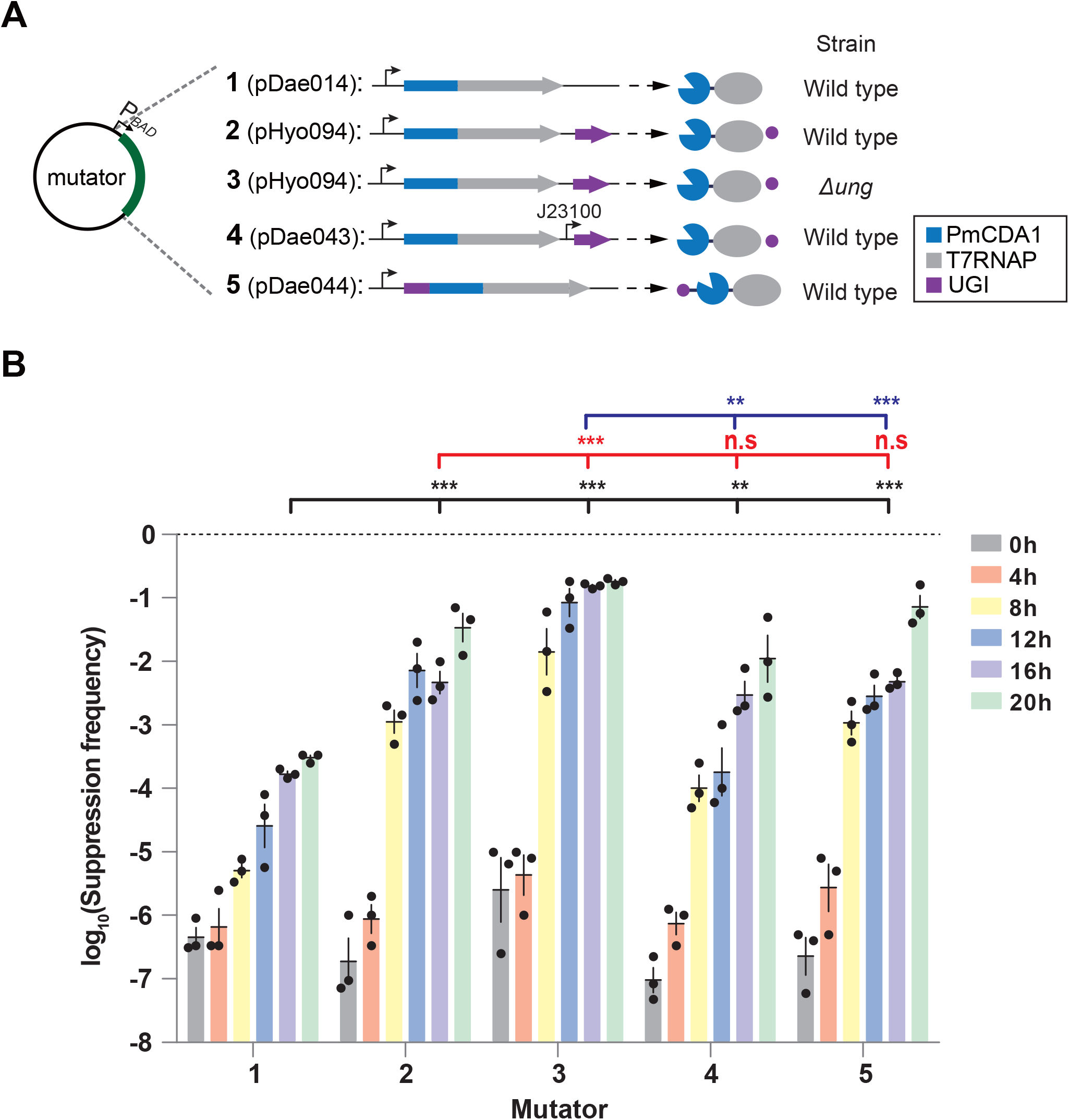
eMutaT7^PmCDA1^ optimization. A Five tested conditions for eMutaT7^PmCDA1^-mediated hypermutation: no UGI in wild-type strain (**1**), UGI expression under the original promoter in wild-type (**2**) or *Δung* (**3**) strains, UGI expression under a new constitutive promoter (J23100) in wild-type strain (**4**), and a triple fusion of UGI, PmCDA1, and T7RNAP in the wild-type strain (**5**). B Frequency of the *pheS*_A294G toxicity suppression at each mutagenesis cycle for cells evolved under the five different conditions shown in (A). Data are presented as dot plots with mean ± standard deviation (SD) (n = 3). \*\**P* < 0.01, \*\*\**P* < 0.001; Student’s *t*-test.

**Figure EV4.**
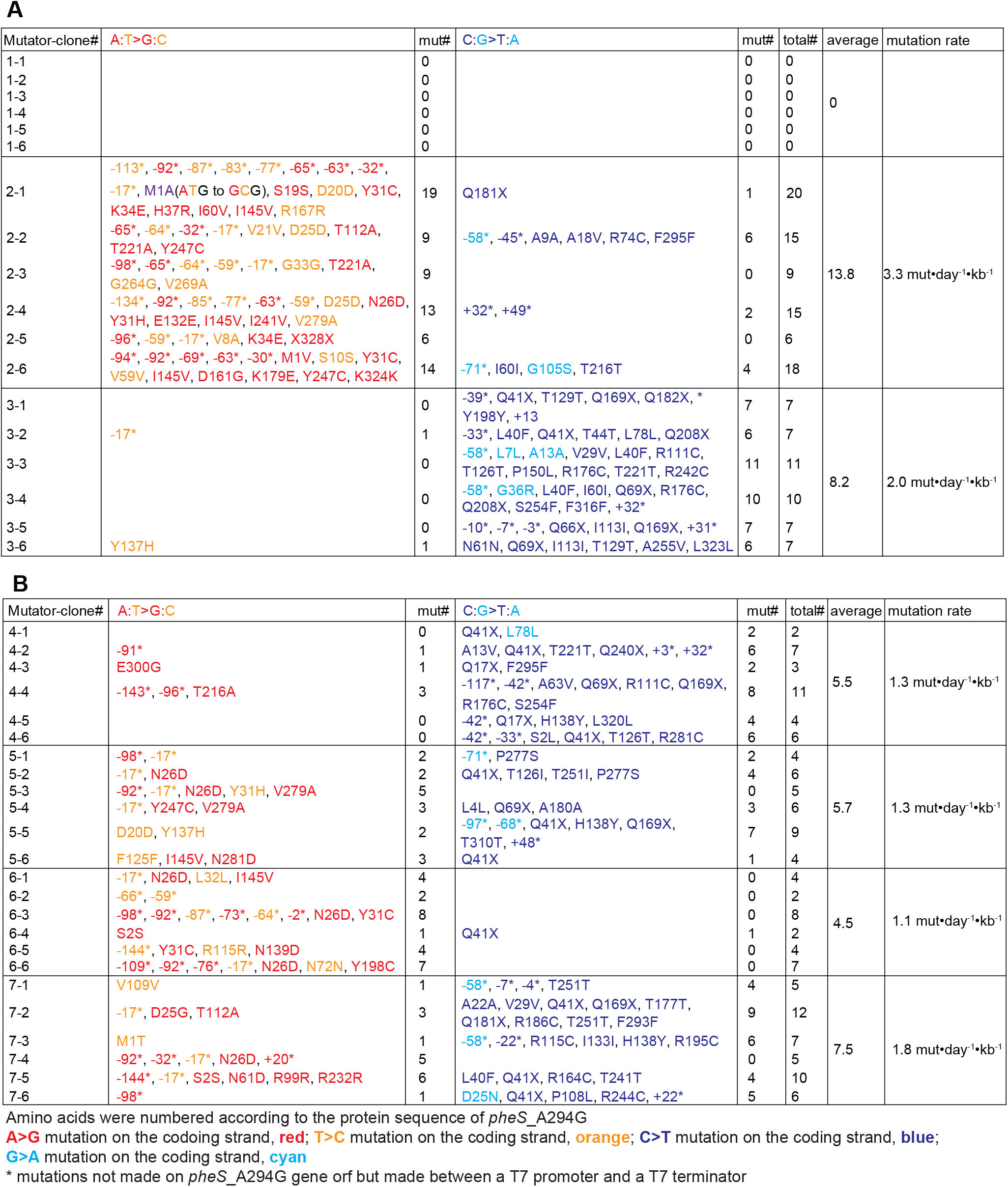
A full list of mutations in samples shown in Figure 4B (A) and 4D (B).

**Figure EV5.**
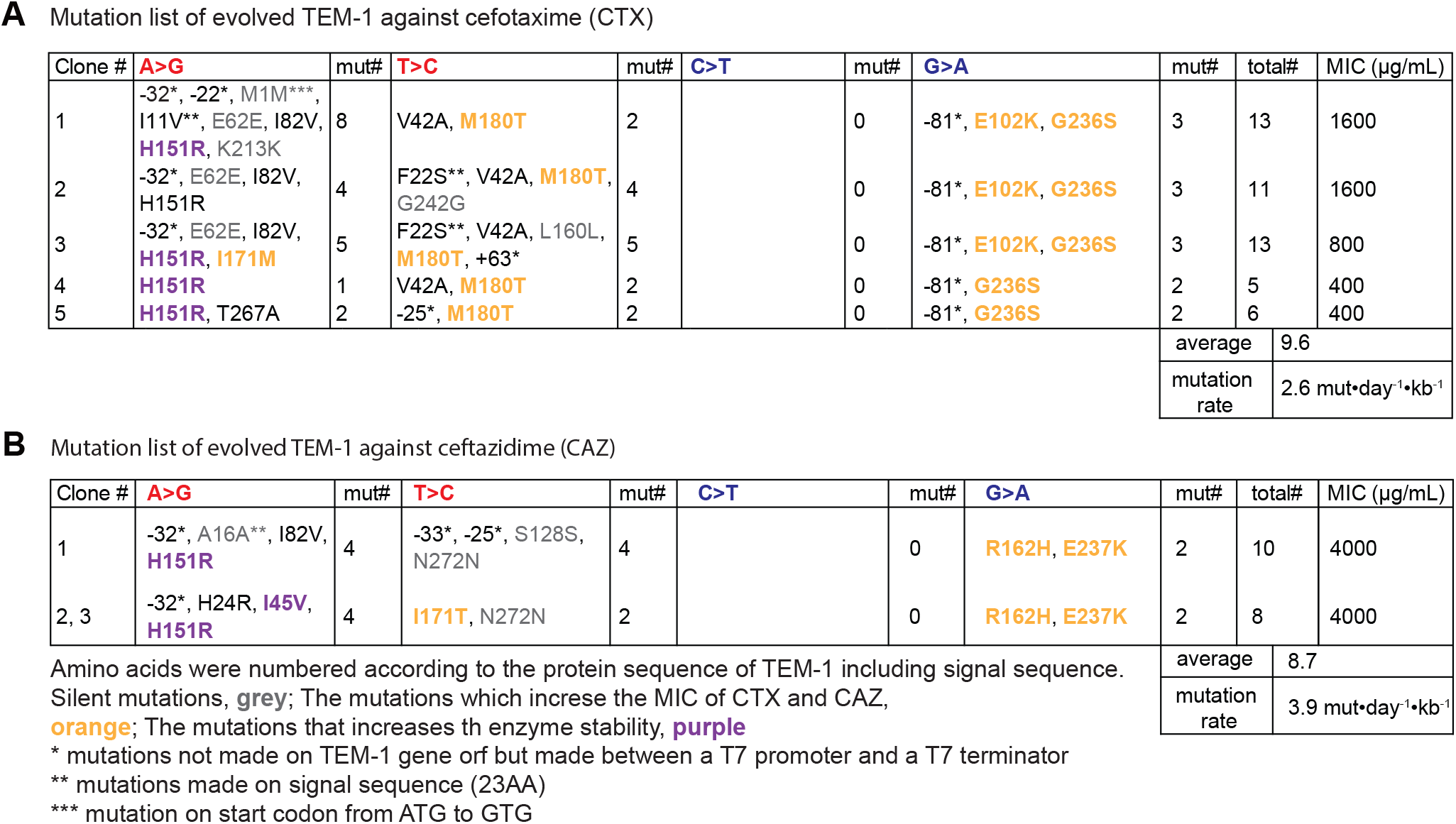
A full list of mutations found in samples shown in Figure 5A (A) and 5B (B).

