## Appendix for "A dual gene-specific mutator system installs all transition mutations at similar rates *in vivo*"

| Contents | Page |
|---|---|
| **Table of Contents** | S1 |
| **Appendix Figure** | S2-S3 |
| **Appendix Tables** | S4-11 |
| **References** | S12 |

**A**

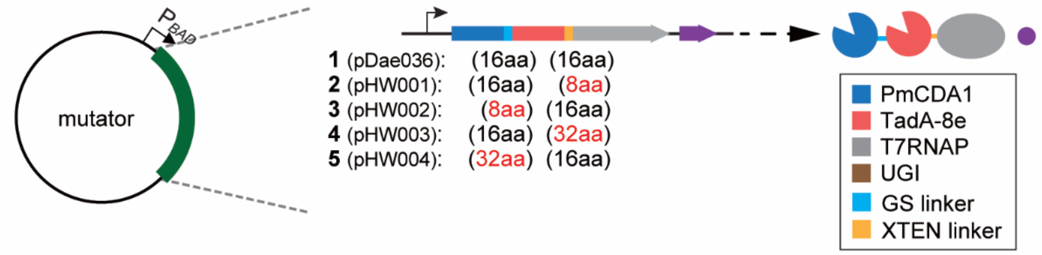


**B**

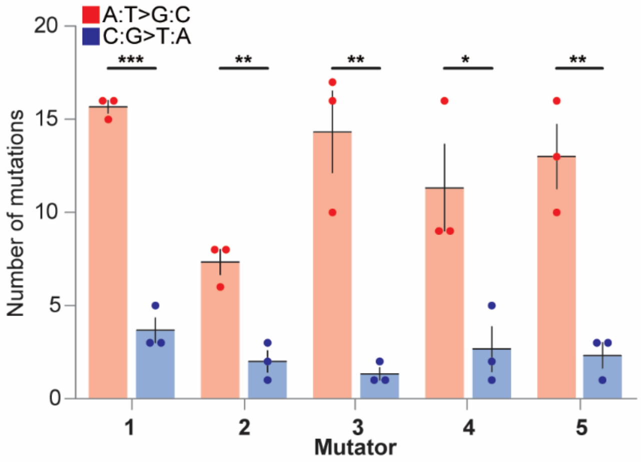


**C**

| Mutator-clone# | A:T>G:C | mut# | C:G>T:A | mut# | total# |
|---|---|---|---|---|---|
| 1-1 | -131*, -111*, -109*, -93*, -92*, -64*, -17*, D25D, Y31H, H148R, T162T, T170A, E210G, T251A, S314S | 15 | -33*, Q41X, T216T | 3 | 18 |
| 1-2 | -98*, -92*, -65*, -64*, -30*, -17*, -1*, E6E, S10G, V21V, D25G(GAT to GGC), Y31R(TAT to CGT), Y198C, K265K | 16 | -71*, F140F, F158F | 3 | 19 |
| 1-3 | -92*, -59*, -17*, S2P, N26D, T39A, M42T, A56A, T112A, D114G, I145V, F160L, V173V, R186C, N233S, K324K | 16 | A5V, I60I, P150S, R186C, P277L | 5 | 21 |
| 2-1 | -92*, -63*, -17*, D25D, V109V, V173V, E236G, +56* | 8 | -70*, -36*, V29V | 3 | 11 |
| 2-2 | -108*, -92*, -59*, G121G, I145T, F220F, +20*, +117* | 8 | A9A, T251T | 2 | 10 |
| 2-3 | -59*, -17*, N61D, L83P, Y137C, +23* | 6 | T251T | 1 | 7 |
| 3-1 | -133*, -93*, -92*, -85*, -66*, -59*, -44*, -17*, N26D, H37R, I145V, T157A, V173A, Q182R, N233S, Y290H | 16 | L4L, T126T | 2 | 18 |
| 3-2 | -98*, -92*, -17*, H3H, I15I, N26D, N72N, F125F, T216A, V269A | 10 | -71* | 1 | 11 |
| 3-3 | -92*, -89*, -82*, -81*, -66*, -64*, -59*, -17*, V8V, S16G, G33G, R74R, N84N, I113V, R167R, V173A, I218V | 17 | G271S | 1 | 18 |
| 4-1 | -131*, -54*, -17*, D20D, R11R, E117E, F125F, T126A, Y137H | 9 | P49L | 1 | 10 |
| 4-2 | -109*, -92*, -77*, -65*, -17*, -6*, -2*, -1*, M1A(ATG to GCG), V21V, D25D, H37R, T91A, Y137C, T168A | 16 | -110*, N61N, R176C, Q181X, P277L | 5 | 21 |
| 4-3 | -131*, -17*, K34E, Y198C, Y247C, T251A, K262E, S314S, +23* | 9 | -71*, L231L | 2 | 11 |
| 5-1 | -98*, -95*, -83*, -82*, -66*, -59*, -17*, L24L, N26N, Y31C, A82A, H138R, I145A(ATT to GCT), T168A, H228R | 16 | Q69X | 1 | 17 |
| 5-2 | -131*, -98*, -96*, -77*, -51*, -17*, G33G, L38L, T112A, T126A, T163A, T177A, I285V | 13 | -68*, Q41X, A58A | 3 | 16 |
| 5-3 | -131*, -98*, -93*, -65*, -60*, -17*, -2*, I145V, H148R, V279A | 10 | -48*, M42I, Q182X | 3 | 13 |

Amino acids were numbered according to the protein sequence of *pheS*_A294G

**A>G** mutation on the codoing strand, **red**; **T>C** mutation on the coding strand, **orange**; **C>T** mutation on the coding strand , **blue**; **G>A** mutation on the coding strand, **cyan**

* mutations not made on *pheS*_A294G gene orf but made between a T7 promoter and a T7 terminator

**Appendix Figure S1. Linker optimization of a triple fusion protein, PmCDA1_TadA-8e_T7RNAP.**

A Design of variants with different linkers: two original linkers (**1**), a shorter linker between TadA-8e and T7RNAP (**2**), a shorter linker between PmCDA1 and TadA-8e (**3**), a longer linker between TadA-8e and T7RNAP (**4**), a longer linker between PmCDA1 and TadA-8e (**5**).

B Number of mutations found in six clones from samples shown in (A) at 20 mutagenesis cycle.

C A list of mutations found in samples shown in (B).

Data are presented as dot plots with mean ± standard deviation (SD) (n = 6). *$P < 0.05$, **$P < 0.01$, ***$P < 0.001$; by Student's *t*-test.

**Appendix Table S1. *E. coli* strains used in this study**

| Strain | Description | Reference |
|---|---|---|
| W3110 | | |
| cHYO057 | W3110 *Δung::kan*$^{R}$ | (Park & Kim, 2021) |
| cDJ085 | W3110 *ΔalkA::Sm*$^{R}$ *Δnfi::Kan*$^{R}$ | This study |

**Appendix Table S2. Plasmids used in this study**

| Plasmid | Construct | Description | reference |
|---|---|---|---|
| pBAD33 | | Experimental control/cloning vector | |
| pHyo094 | pBAD33-PmCDA1-T7RNAP, ugi | eMutaT7$^{PmCDA1}$, source of PmCDA1 | (Park & Kim, 2021) |
| pHyo182 | pVS133-lacI, pheS_A294G | Target plasmid (pheS_A294G) | (Park & Kim, 2021) |
| pHyo183 | pBAD33-T7RNAP, ugi | control | (Park & Kim, 2021) |
| pHyo245 | pVS133-dualT7_pheS_A294G | Dual promoter system/cloning vector | (Park & Kim, 2021) |
| pREMCM3 | | Tetracycline resistance gene | (Melancon & Schultz, 2009) |
| TadA_mut | BamH1-GS-TadA*-GS-EcoR1 | Source of TadA-7.10 gene | (Gaudelli *et al*, 2017) |
| pDae027 | pCDF-TadA-8e-TadA-8e | Source of TadA-8e gene | (Richter *et al*, 2020) |
| pDae014 | pBAD33-PmCDA1-T7RNAP | eMutaT7$^{PmCDA1}$ without *ugi* | This study |
| pDae028 | pBAD33-TadA-7.10-T7RNAP | eMutaT7$^{TadA\text{-}7.10}$ | This study |
| pDae029 | pBAD33-TadA-8e-T7RNAP | eMutaT7$^{TadA\text{-}8e}$ | This study |
| pDae032 | pBAD33-TadA-8e | Control | This study |
| pDae035 | pBAD33-TadA-8e-T7RNAP, PmCDA1-T7RNAP, ugi | the expression of two mutators, eMutaT7$^{TadA\text{-}8e}$ and eMutaT7$^{PmCDA1}$ | This study |
| pDae036 | pBAD33-PmCDA1-TadA-8e-T7RNAP, ugi | triply-fused protein of PmCDA1, TadA-8e, and T7RNAP | This study |
| pDae037 | pBAD33-TadA-8e-PmCDA1-T7RNAP, ugi | triply-fused protein of TadA-8e, PmCDA1 and T7RNAP | This study |
| pDae043 | pBAD33-PmCDA1-T7RNAP, J23100(promoter), ugi | Constitutive promoter for *ugi* | This study |
| pDae044 | pBAD33-ugi-PmCDA1-T7RNAP | triply-fused protein of *ugi*, PmCDA1, and T7RNAP | This study |

| | | | |
|---|---|---|---|
| pDae069 | pBAD33-PmCDA1-T7RNAP, (RBS) ugi | eMutaT7$^{PmCDA1}$ with optimized RBS | This study |
| pDae079 | pBAD33-TadA-8e-T7RNAP, PmCDA1-T7RNAP, (RBS) ugi | eMutaT7$^{transition}$ | This study |
| pDae080 | pBAD33-PmCDA1-T7RNAP, (RBS) ugi, TadA-8e-T7RNAP | the expression of two mutators, eMutaT7$^{PmCDA1}$ and eMutaT7$^{TadA\text{-}8e}$ and optimized *ugi* | This study |
| pHW001 | pBAD33-PmCDA1-16aa-TadA-8e-8aa-T7RNAP, ugi | Shoter XTEN linker | This study |
| pHW002 | pBAD33-PmCDA1-8aa-TadA-8e-16aa-T7RNAP, ugi | Shoter GS linker | This study |
| pHW003 | pBAD33-PmCDA1-16aa-TadA-8e-32aa-T7RNAP, ugi | Longer XTEN linker | This study |
| pHW004 | pBAD33-PmCDA1-32aa-TadA-8e-16aa-T7RNAP, ugi | Longer XTEN linker | This study |
| pGE158 | pVS133-dualT7_ss-TEM-1, tetR | Evolution target (TEM-1) | This study |

**Appendix Table S3. Primers used in this study**

| Oligonucleotides | Sequence (5'→3') | Description |
|---|---|---|
| T7promoter | TAATACGACTCACTATAGGG | Universal sequencing primer |
| T7terminator | GCTAGTTATTGCTCAGCGG | Universal sequencing primer |
| pBAD-F | ATGCCATAGCATTTTTATCCA | Universal sequencing primer |
| pBAD-R | GATTTAATCTGTATCAGG | Universal sequencing primer |
| 022_PxUgT_PxT_ovlp_fw | aacacgattaacatcgctaagaacg | Cloning of TadA-8e-T7RNAP(pDae029) |
| 029_pBAD_Gibson_rv | CCATGGtgaattcctcctGagctcg | Cloning of TadA-7.10-T7RNAP(pDae028) |
| 052_T7RNAPQ265_rv | gttgacgctcaaacatcttgc | |
| 053_PmCDA_P57_rv | gggcttgttgacggcatag | |
| 054_PmCDA_G137_fw | ggactctggaatctgaggg | |
| 055_UPT-GS_ovlp_fw | GTAGCGGCTCTGGTTCCGGCTCTGGTAGCGGATCCAcagacgccgagtacgtg | Clonging of pBAD33-TadA-8e-PmCDA1-T7RNAP, ugi (pDae037) |
| 058_PUT2_GS_ovlp_rv | CCGGAACCAGAGCCGCTACCAGAGCCGGAACCaacggctggagacttagtgg | Clonging of pBAD33-PmCDA1-TadA-8e-T7RNAP, ugi (pDae036) |
| 062_pYH103_ovlp1_rv | ATGCCATGGtgaattcctc | Clonging of pBAD33-TadA-8e-PmCDA1-T7RNAP, ugi (pDae037) |
| 063_PmCDA_ovlp1_fw | ctCaggaggaattcaCCATGGC | Clonging of pBAD33-TadA-8e-PmCDA1-T7RNAP, ugi (pDae037) |
| 128_pYH103_dpolI_rv | catATGCCATGGtgaattcctc | Cloning of TadA-8e-T7RNAP(pDae029) |
| 129_nfi_fw | GGTCACGGCATTCATCAGG | |
| 130_nfi_rv | GACATGCTGCCAGCTTTCC | |
| 131_alkA_fw | GCGAAATGTTGCCGTCGC | |
| 132_alkA_rv | CCCATCGCCTGATGCGAC | |
| 133_TadA8e_IVA_fw | ctCaggaggaattcaCCATGGCATATGAGTGAAGTTGAATTCAGCCATG | Cloning of TadA-8e-T7RNAP(pDae029) |
| 134_TadA8e_IVA_rv | gaagtcgttcttagcgatgttaatcgtgttactttcgggtgtggcg | Cloning of TadA-8e-T7RNAP(pDae029) |
| 137_TadAdimer_del_fw | GGTTCCGGTAGCTTGTCTGAAGTC | Cloning of TadA-7.10-T7RNAP(pDae028) |
| 138_TadA_del_fw | CATATGTCTGAAGTCGAATTTAGCCACG | Cloning of TadA-7.10-T7RNAP(pDae028) |
| 161_dalkA_SmR3_fw | ATGGCGGCAAAATTGACCGCCAGAGTGGCACAGCTTTATGgcgaccgagtgagctagctatttg | Amplification of streptomycin resistance gene from pCDF for construction of alkA k/o strain |
| 162_dalkA_SmR3_rv | GGGAAGCAGATATACTCCGGAAAATCATCCAGCCGTTCGCgaacgaattgttagacattatttgcc | Amplification of streptomycin resistance gene from pCDF for construction of alkA k/o strain |

| | | |
|---|---|---|
| 163_Am7(8e)_dL_rv | ggagtctcgctgccgcttaATTAATG CTG | Clonging of TadA-8e(pDae032) |
| 165_Am7(8e)_dT7_fw | ggacttcgcgttcgcgtaa | Clonging of TadA-8e(pDae032) |
| 170_duet_Am7_IVA2_fw | cagaatttgcctggcggcagacttttcata ctcccgccattcagagaag | Clonging of pDae035, pDae079 (=eMutaT7$^{transition}$), and pDae080 |
| 171_duet_tm_Am7_IVA1_rv | cagggttattgtctcatgagcg | Clonging of pDae035, pDae079 (=eMutaT7$^{transition}$), and pDae080 |
| 172_duet_PxT_IVA2_rv | ctgccgccaggcaaattc | Clonging of pDae035, pDae079 (=eMutaT7$^{transition}$), and pDae080 |
| 173_duet_tm_PxT_IVA1_fw | gtatccgctcatgagacaataacc | Clonging of pDae035, pDae079 (=eMutaT7$^{transition}$), and pDae080 |
| 174_PgAxT_PxT_IVA1_fw | ctcccgggacctcagagtc | Clonging of pBAD33-PmCDA1-TadA-8e-T7RNAP, ugi (pDae036) |
| 175_PgAxT_TadA8_IVA1_fw | GTAGCGGCTCTGGTTCCGG CTCTGGTAGCGGATCCAGT GAAGTTGAATTCAGCCATG | Clonging of pBAD33-PmCDA1-TadA-8e-T7RNAP, ugi (pDae036) |
| 176_PgAxT_TadA8_IVA1_rv | ggactctgaggtcccggg | Clonging of pBAD33-PmCDA1-TadA-8e-T7RNAP, ugi (pDae036) |
| 177_AgPxT_TadA8_IVA2_rv | CCGGAACCAGAGCCGCTAC CAGAGCCGGAACCATTAAT GCTGCTCTGTGCTTTCT | Clonging of pBAD33-TadA-8e-PmCDA1-T7RNAP, ugi (pDae037) |
| 182_dnfi_KanR_fw | ATGGATCTCGCGTCATTACG CGCTCAACAAATCGAACTG GCTTGATCCTTTGATCTTTT CTACGGGGtc | Amplification of kanamycin resistance gene from pET28b for construction of nfi k/o strain |
| 183_dnfi_KanR_rv | TTAGGGCTGATTTGCTGTAT AGCGCACGAACGCCGGACG TTCCGATGGCACTTTTCGG GGAAATGTG | Amplification of kanamycin resistance gene from pET28b for construction of nfi k/o strain |
| 184_ugi-F | caaaccctgggctctggtg | |
| 185_Amp-R | cagcatcttttactttcaccagc | |
| 186_T7RNAP_PstI_rv | atactgcagttacgcgaacgcgaagtcc | Clonging of pBAD33-PmCDA1-T7RNAP (pDae014) |
| 192_ugi_IVA-F | ctCaggaggaattcaCCATGGCAT atgaccaacctttccgacatc | Cloning of pBAD33-ugi-PmCDA1-T7RNAP (pDae044) |
| 193_ugi_IVA-R | CCGGAACCAGAGCCGCTAC CAGAGCCGGAACCtagcatcttg atcttgttctctcc | Cloning of pBAD33-ugi-PmCDA1-T7RNAP (pDae044) |

| | | |
|---|---|---|
| 194_ugi-J23100-R | ctaggactgagctagccgtcaaaggatc cccgggctgcaG | Cloning of pBAD33-PmCDA1-T7RNAP, J23100(promoter), ugi (pDae043) |
| 195_ugi-J23100-RBS-F | gtacagtgctagcctagagtcaggagg agacctgcgatgaccaacctttcc | Cloning of pBAD33-PmCDA1-T7RNAP, J23100(promoter), ugi (pDae043) |
| 223-pHyo250-IVA-F | AGCACCACCACCACCACCA CTG | Cloning of pVS133-dualT7_ss-TEM-1, tetR (pGE158) |
| 224-pHyo250-IVA-R | CATGGTATATCTCCTTCTTA AAGTTAAACAAAA | Cloning of pVS133-dualT7_ss-TEM-1, tetR (pGE158) |
| 225-TEM1-IVA-F | CCTCTAGAAATAATTTTGTTT AACTTTAAGAAGGAGATATA CCatgagtattcaacatttccgtgtcg | Cloning of pVS133-dualT7_ss-TEM-1, tetR (pGE158) |
| 226-TEM1-IVA-R | CAGTGGTGGTGGTGGTGGT GCTctcgaGttaccaatgcttaatcagt gagg | Cloning of pVS133-dualT7_ss-TEM-1, tetR (pGE158) |
| 231-AxT-32aa-F | CTCTGGTTCCGGCTCTGGT AGCGGATCCAgcggcagcgaga ctccc | Cloning of pBAD33-PmCDA1-16aa-TadA-8e-32aa-T7RNAP, ugi (pHW003) |
| 232-AxT-32aa-R | CCGCTACCAGAGCCGGAAC CATTAATGCTGCTCTGTGCT TTCTTTTGTG | Cloning of pBAD33-PmCDA1-16aa-TadA-8e-32aa-T7RNAP, ugi (pHW003) |
| 235-GS-32aa-F | ctcagagtccgccacacccgaaagtA GTGAAGTTGAATTCAGCCAT G | Cloning of pBAD33-PmCDA1-32aa-TadA-8e-16aa-T7RNAP, ugi (pHW004) |
| 236-GS-32aa-R | gtcccgggagtctcgctgccAcTGGA TCCGCTACCA | Cloning of pBAD33-PmCDA1-32aa-TadA-8e-16aa-T7RNAP, ugi (pHW004) |
| 239_PstI_pBAD_fw | ataaCTGCAGgcatgcaagc | Clonging of pBAD33-TadA-7.10-T7RNAP |
| 240-Ftet-IVA-F | ctgtcagaccaagtttactcaactg | Cloning of pVS133-dualT7_ss-TEM-1, tetR (pGE158) |
| 241-Ftet-IVA-R | gacataagtccatcagttcaacgg | Cloning of pVS133-dualT7_ss-TEM-1, tetR (pGE158) |
| 242-TcR-IVA-F | gacttccgttgaactgatggacttatgtcg taattctcatgtttgacagcttatcatc | Cloning of pVS133-dualT7_ss-TEM-1, tetR (pGE158) |
| 243-TcR-IVA-R | cgcagttgagtaaacttggtctgacagtg gagtggtgaatccgttagc | Cloning of pVS133-dualT7_ss-TEM-1, tetR (pGE158) |

| | | |
|---|---|---|
| 245-G8aa-F | ccgaaagtAGTGAAGTTGAATTCAGCCATGAATATTG | Cloning of pBAD33-PmCDA1-8aa-TadA-8e-16aa-T7RNAP, ugi (pHW002) |
| 246-G8aa-R | gtgtggcggactctgaaacggctggagacttagtgg | Cloning of pBAD33-PmCDA1-8aa-TadA-8e-16aa-T7RNAP, ugi (pHW002) |
| 247-X8aa-F | CTCTGGTAGCGGCTCTaacacgattaacatcgctaagaacg | Cloning of pBAD33-PmCDA1-16aa-TadA-8e-8aa-T7RNAP, ugi (pHW001) |
| 248-X8aa-R | CCGGAACCATTAATGCTGCTCTGTGCTTTCTTTTGTG | Cloning of pBAD33-PmCDA1-16aa-TadA-8e-8aa-T7RNAP, ugi (pHW001) |
| 254_pBAD-NdeI_rv | aatCATATGCCATGGtgaattcctc | Clonging of pBAD33-TadA-7.10-T7RNAP |
| 255-TadA-A113-R | GCACCGCGCTTGCTATTAC | |
| 314-F-seq-F2 | CATTAGGAAGCAGCCCAGTAGTAG | sequencing primer for target gene |
| 315-F-seq-R2 | gagacgaaagggcccgtacg | sequencing primer for target gene |
| 316-UGI-RBS-F | CAATAAATAAGGAGGATTTTTTatgaccaacctttccgacatcatagag | Cloning of pBAD33-PmCDA1-T7RNAP, (RBS) ugi (pDae069) |
| 318-UGI-RBS-R | CTAGTACTCAAACAGAGCGCGCTCTGTTaggatccccgggctgcaG | Cloning of pBAD33-PmCDA1-T7RNAP, (RBS) ugi (pDae069) |
| 341_NdeI_tadA_fw | TTAcatatgTTGTCTGAAGTCGAATTTAGCCAC | Clonging of pBAD33-TadA-7.10-T7RNAP |
| 367_T7RNAP_taa_PstI_rv | ttatctgcagttacgcgaacgcgaagtcc | Clonging of pBAD33-TadA-7.10-T7RNAP |
| 368_28b_th_rv | CATATGGCTGCCGCG | Clonging of pET28b-TadA-7.10-T7RNAP |
| 369_28b_ov_tadA_fw | GGTGCCGCGCGGCAGCCATATGTTGTCTGAAGTCGAATTTAGCCACG | Clonging of pET28b-TadA-7.10-T7RNAP |
| 370_tadA_XTEN_ov_rv | gtggcggactctgaggtcccgggagtctcgctgccgctATCCGTCGAGGATTGCG | Clonging of pET28b-TadA-7.10-T7RNAP |
| 371_ov_XTEN_T7RNAP_fw | gactcccgggacctcagagtccgccacacccgaaagtaacacgattaacatcgctaagaac | Clonging of pET28b-TadA-7.10-T7RNAP |

| | | |
|---|---|---|
| 473_pBAD_dugi_fw__w187 | aagcttggctgttttggc | Clonging of pBAD33-PmCDA1-T7RNAP (pDae014) |
